# Descending neurons integrate learnt information from mushroom body with context to promote escape behaviour

**DOI:** 10.1101/2025.09.30.679458

**Authors:** Benjamin M. W. Jones, Samuel N. Harris, Nicolo G. Ceffa, Albert Cardona, Marta Zlatic

## Abstract

To behave adaptively in the environment and select appropriate responses to sensory stimuli, animals integrate learnt information about the valences of stimuli with contextual information. While significant progress has been made in understanding how animals learn which stimuli predict positive and negative outcomes, how the outputs of associative learning circuits flexibly promote different behaviours, depending on context, is not well understood. Addressing this question requires mapping the pathways from learning circuits to nerve cord command neurons that promote specific actions, and understanding where and how contextual information converges with these pathways to modulate their activity. These are daunting tasks in larger brains where synaptic-resolution connectivity maps are not available. We, therefore, addressed this question in the tractable *Drosophila* larva using a combination of connectomic analyses, imaging and manipulation of neural activity and behavioural analysis. We characterised, with synaptic-resolution, the pathways from the higher-order learning circuit (the mushroom body, MB) all the way to a specific cord command neuron (Goro) in the nerve cord that promotes the most vigorous escape response, rolling. Rolling is the fastest, but also the most energetically costly escape response and is activated by multisensory cues in the context of predator attack. We identify a pair of brain descending neurons, Ipsigoro, that integrate learnt information (via input from MBONs) and nociceptive context (via input from ascending neurons) to facilitate rolling via direct inputs to Goro. Our study reveals the circuit mechanism by which context and learnt information are integrated by brain descending neurons to activate specific nerve cord command neurons and promote specific actions.

## Introduction

To behave adaptively in the environment and select appropriate responses to sensory stimuli, animals integrate learnt information about the valences of stimuli with contextual information. For example, if an animal has learnt to associate an odour with a predator, it could avoid that odour in different ways, depending on context: if it senses the odour without other signs of predator, it may slowly move away, but if it senses the odour and hears a noise it may run away. While significant progress has been made in recent years in understanding the neural circuits involved in associative learning, how the outputs of these circuits flexibly promote different avoidance (or approach) behaviours, depending on context is not well understood.

Brain structures such as the Striatum (vertebrates) and the Mushroom Body (MB) (invertebrates) carry out classical conditioning (Eichler et al., 2017; Heisenberg, 2003; Suri & Schultz, 1999). This process allows animals to associate conditioned stimuli (CSs) with rewards or punishments (Pavlov, 1927). The output of these structures represents the stimulus’ learned value which can influence the selected behavioural response (Aso, Sitaraman, et al., 2014; Costa et al., 2019; Eschbach et al., 2021). Specific behavioural outcomes are determined by specific motor command or command-like neurons (Jing, 2009; Kupfermann & Weiss, 1978). They represent a bottleneck for all the information contributing to action selection including both learned and innate drives (Sokolov & Nezlina, 2008). As such, the influence of a CS on behaviour will depend on its learned value and the innate context (Devineni & Scaplen, 2022; McDannald, 2023). The circuits mechanisms by which learned value signals and innate contextuals are integrated to activate specific command neurons in the nerve cord and trigger specific actions are not well understood.

Addressing these questions requires mapping the neural pathways between the learning circuit and specific nerve cord command neurons and investigating how contextual information converges with this circuitry, which is extremely challenging in larger model systems. We therefore use the tractable *Drosophila* larva to investigate this problem, which has a rich behavioural repertoire (Eschbach & Zlatic, 2020; Ohyama et al., 2013) and identified command neurons (Clark et al., 2018; Ohyama et al., 2015; Takagi et al., 2017), extensive genetic toolkit (Jenett et al., 2012; Kvon et al., 2014; Lai & Lee, 2006; Owald et al., 2015) and fully reconstructed brain connectome (Winding et al., 2024) which includes the MB (Eichler et al., 2017).

Using this tractable model-system, we investigated the neural circuit mechanism by which learnt information and context are integrated to select the most vigorous escape behaviour, rolling. Across the animal kingdom, escape behaviours are only triggered in the presence of specific forms of aversive stimulation (Branco & Redgrave, 2020; Tracey et al. 2003; Ohyama et al. 2015). While these behaviours are often considered hardwired to these aversive inputs, there is a cost associated with both withholding and triggering them inappropriately. If the behaviour is not triggered when needed, the animal may face predation. If the behaviour is triggered when not required, the animal may flee a resource rich environment unnecessarily, expending large amounts of energy in the process. It is more adaptive to dynamically modulate the threshold for escape (Ydenberg & Dill, 1986). For example, the likelihood of selecting an escape has been shown to be modulated by habituation (Zucker, 1972), sensitisation (Krasne & Glanzman, 1986), prior or concurrent food consumption (Advokat, 1980; Casey & Morrow, 1983; Krasne & Lee, 1988) and the presence of other stimuli such as food odour (Liden et al., 2010), social (Domenici & Batty, 1997), or mechanosensory cues (Ohyama et al. 2015).

For *Drosophila* larvae, rolling is the fastest, but also the most energetically costly escape response, activated, primarily, by multisensory cues in the context of predator attack. We have previously identified a set of command neurons in the nerve cord (called Goro), whose activation is sufficient to trigger rolling. Goro receives convergent input from nociceptive circuits in the nerve cord, but also from brain descending neurons. Here, we demonstrate that an aversively conditioned odour promotes rolling, in a context-dependent manner. We identify a functional, excitatory descending input to the command neuron for rolling escape and demonstrate that this neuron receives input from both the MB and innate nociceptive pathways. Together, our study reveals a mechanism for context-dependent memory-based action selection.

## Results

### Nociceptive context modulates the selection of larval escape response

First, we asked whether contextual information is integrated with learnt information to bias the selection of larval escape response. When larvae are trained by repeatedly pairing an odour (CS^+^) with a noxious cue (US), they learn to avoid the CS. Noxious stimulation alone induces rolling in a significant fraction of animals, but even though larvae learn that the CS^+^ predicts the noxious cue, when presented with CS^+^ alone, after training, they do not roll. Instead, they avoid the CS by turning and crawling away. We wondered whether presenting US together with CS, during testing, will increase the likelihood of rolling, compared to US alone.

We used a combination of optogenetic activation of Or42b olfactory neurons (as a CS) and thermogenetic activation of primary nociceptive interneurons, called Basins (as a US), to establish aversive memory, as previously described (Croteau-Chonka et al., 2022). Two groups of larvae were used for the experiments which received different stimulation protocols. In the forward-paired group, the CS^+^ was predictive of the US during training while in the backward-paired group, the CS^-^ was presented after the US and was therefore not predictive of punishment.

To investigate the ability of larvae to express a Basin-associated memory in a context-dependent manner, Basin punishment was present during the testing period. The full training and testing protocol is displayed in **Figure 1a**. The two groups of animals received 24 training trials on an agarose plate. Animal outlines were captured by an overhead camera and rolling during testing was manually annotated. We found that the forward-paired group spent significantly more time rolling in response to the combination of CS and US (**Figures 1b-c**), than those in the backward-paired group indicating that nociceptive context and learnt information are integrated to increase the likelihood of rolling escape. Specifically, more rolling behaviour is seen in the presence of a CS^+^ which is predictive of Basin punishment when compared to an CS^-^ which has been presented the same number of times but not paired with punishment. This is similar to the fear-potentiated startle paradigm in vertebrates where a CS can both drive freezing when presented in isolation and potentiate startle in the presence of an auditory stimulus (Brown et al., 1951; M. Davis et al., 1993).

**Figure 1.**
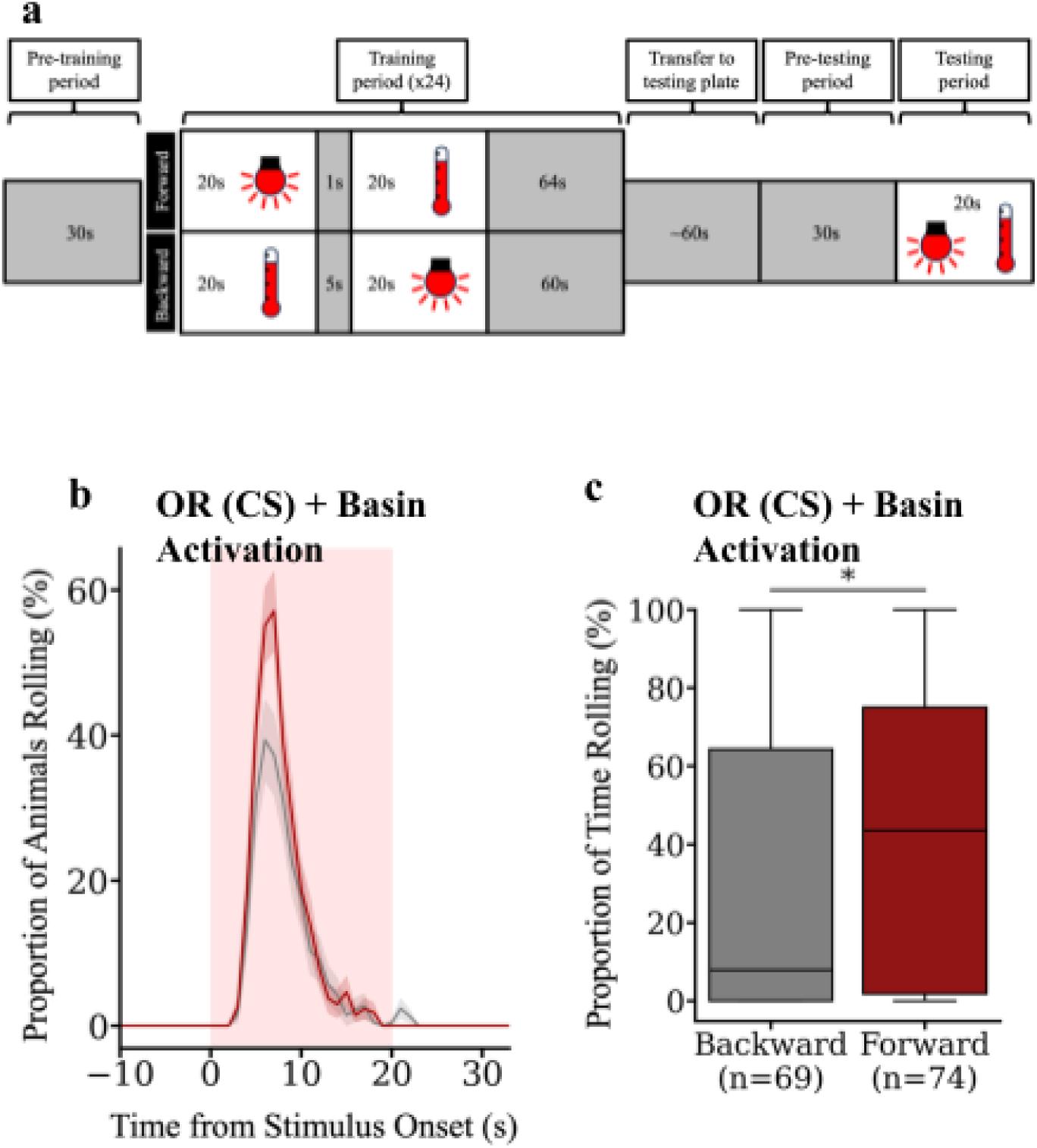
Aversively conditioned odour facilitates rolling in nociceptive context. **a)** Schematic of the protocol for aversive classical conditioning. After an initial pre-training period of 30s, 24 training trials were delivered. The odour CS was optogenetic activation of Or42b neurons with red-light. The punishment (unconditioned stimulus, US) was thermogenetic activation of Basin neurons with an IR laser. Stimulation was delivered at the start of each 105s trial. Both stimuli were delivered for 20s. In the forward-paired group odour preceded punishment separated by a 1s gap. In the backward-paired group, odour followed the offset of punishment with a 5s gap. For the remainder of the trial there was no stimulation. At the end of training, larvae were transferred to a testing agarose plate. After a 30s pre-testing period, larvae received 20s of simultaneous red-light and heat. **b)** Time-series plot of the percentage of animals rolling during the testing period. Data were down-sampled to 1Hz to improve visualisation. Red shading indicates the delivery of the light and heat stimulus. Error shading shows the mean rolling ± s.e.m. (Or42b-lexA>lexAop-Chrimson, Basin-Gal4>UAS-CsChrimson)**. c)** Boxplot of the percentage of time animals spent rolling during testing. Analysis was restricted to a 5s window starting at 4s after the start of stimulation. Boxes show medians and upper and lower quartiles. Whiskers show the range of data. The forward-paired group spent significantly longer rolling than the backward-paired group. Statistics were calculated with two-sided Mann-Whitney U test; * p<0.05.

### The roll-command neuron, Goro, receives mushroom body input via a descending neuron, Ipsigoro

Classical odour conditioning in *Drosophila* adult and larva is mediated by the MB (Heisenberg et al., 1985; Eschbach et al. 2020), located in the brain. In contrast, the rolling command neuron, Goro, and the nociceptive Basin interneurons are located in the nerve cord (Ohyama et al., 2015; Takagi et al., 2017; Burgos et al., 2018). Therefore, for an aversive CS to influence nociceptive rolling behaviour as shown in **Figure 1b-c**, information from the brain must be transmitted to roll-evoking nerve cord command neurons. We, therefore, wanted to identify a functional descending input to Goro which receives structural pathways from the MB and can modulate rolling.

Synaptic inputs to Goro have been previously reconstructed in an EM volume spanning the entire larval nervous system and two descending neurons (DNs), named Ipsigoro and Contragoro, have been identified which form synapses onto Goro (Ohyama et al., 2015). The recent reconstruction of the larval brain connectome allowed us to investigate the inputs to these DNs, including those from the MB (Eichler et al., 2017; Winding et al., 2024).

While Ipsigoro and Contragoro have been shown to synapse onto ipsilateral and contralateral Goro neurons, respectively (Ohyama et al., 2015), we first wanted to check whether these connections are strong and reproducible [i.e. whether they “account for ≥1% input” onto Goro and whether they are “observed between homologous pre-and postsynaptic partners in both brain hemispheres” (Winding et al., 2024)]. The strongest connections are from Ipsigoro to the ipsilateral Goro neuron, with an average input percentage of 2.4% (**Figure 2a**). The input from Contragoro is weaker and accounts for only 0.6% of Goro input. This suggests that Ipsigoro is the dominant DN input to Goro and is therefore a particularly good candidate for its descending control.

**Figure 2.**
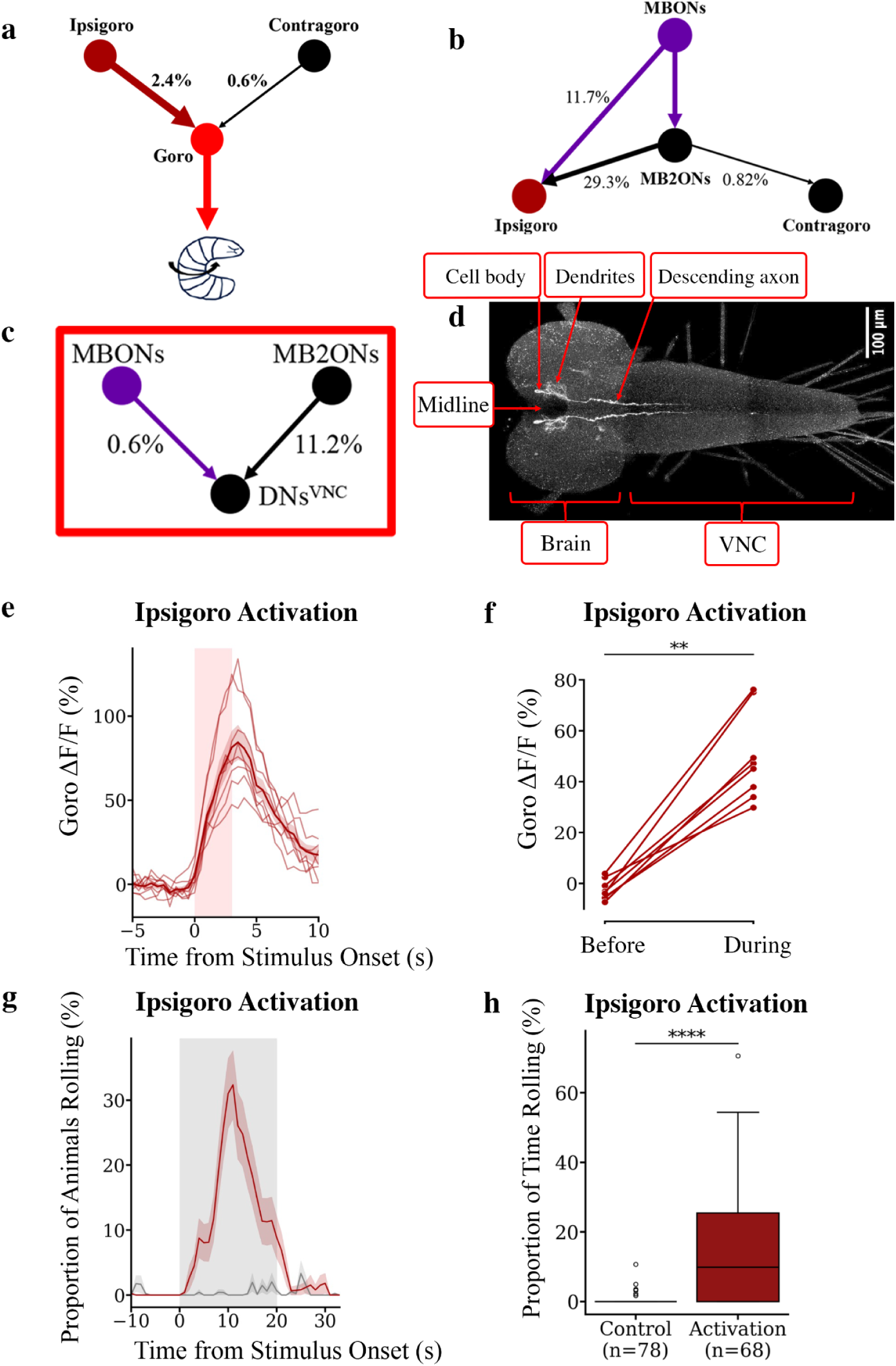
A descending neuron (Ipsigoro), downstream of the MB activates Goro and facilitates rolling. a-c) Structural connectivity suggests Ipsigoro could mediate MB-Goro interactions. **a)** The percentage input onto Goro neurons from ipsilateral and contralateral pathways originating from Ipsigoro and Contragoro respectively. Left and right homologous neurons are merged into single nodes with the mean input percentage displayed. Only the pathway from Ipsigoro meets the ≥1% threshold for a strong connection suggesting it is the dominant descending synaptic input onto Goro. **b)** Percentage contribution of all synapses onto Ipsigoro and Contragoro originating from mushroom body output neurons (MBONs) and their strong downstream partners (MB2ONs). Contragoro receives little input from these neurons while the Ipsigoro pair receives over 40% of its input synapses from them. This suggests that Ipsigoro is the best candidate for mediating MB-Goro interactions. **c)** Percentage contribution of all synapses onto all other brain neurons descending to the VNC (DNs^VNC^) originating from MBONs and MB2ONs. The percentages are considerably lower than for Ipsigoro with around a 12% contribution from these inputs suggesting Ipsigoro is specialised to integrate information from the MB. **d) A Split-GAL4 line (Ipsigoro-Gal4) selectively drives expression in Ipsigoro.** Max intensity projection of a 3D light microscopy volume showing selective expression of UAS-*myr*-GFP in the pair of Ipsigoro descending neurons. **e-h) Ipsigoro can activate Goro and evoke rolling escape**. **e)** Time-series plot of Goro ΔF/F averaged over three windows of optogenetic Ipsigoro activation and over three recordings (nine total stimulations). Thin lines show data for each individual animal, the thick line shows the mean of all animals. Data was downsampled to 2Hz to improve visualisation. Red shading indicates the delivery of the red-light stimulus. Error shading shows the mean ± s.e.m. An excitatory response in Goro is seen during Ipsigoro stimulation (Ipsigoro-Gal4>UAS-CsChrimson, Goro-lexA>lexAop-GCaMP6s). **f)** Mean Goro ΔF/F is significantly higher during Ipsigoro stimulation as compared to the 3s prior suggesting the presence of an excitatory connection. Statistics were conducted using a Wilcoxon signed-rank test; ** p<0.01, N=8. **g)** Time-series plot of the percentage of animals rolling with or without thermogenetic activation of Ipsigoro. Data were down-sampled to 1Hz to improve visualisation. Grey shading indicates the delivery of the heat stimulus. Error shading shows the mean rolling ± s.e.m. ((Ipsigoro-Gal4 or empty-Split-Gal4)>UAS-dTrpA1). **h)** Boxplot of the percentage of time animals spent rolling during the 20s stimulation window. Boxes show medians and upper and lower quartiles. Whiskers show the upper quartile + 1.5*IQR with outliers plotted as empty circles. There was significantly more heat-evoked rolling in the Ipsigoro activation group suggesting this neuron facilitates rolling escape. Statistics were calculated with a two-sided Mann-Whitney U test; **** p<0.0001.

Next, we investigated what proportion of input to the DNs comes from the MB, to determine how likely they are to encode learned value (**Figure 2b**). We examined the percentage of all synapses onto the two DNs that came from MB output neurons (MBONs) or their strongly connected downstream partners (MB2ONs) (Eichler et al., 2017; Eschbach et al., 2021). Contragoro neurons receive no input from MBONs and only very limited input from MB2ONs (**Figure 2b**). Conversely, Ipsigoro neurons receive approximately 42% of their synapses from the two classes of neuron with approximately 12% of inputs coming directly from MBONs (**Figure 2b**). To determine how typical this is of DNs, we examined the percentage contribution of MBONs and MB2ONs to the pooled synaptic inputs to all brain DNs, excluding Ipsigoro and Contragoro (Winding et al., 2024). **Figure 2c** shows that these neurons receive a much smaller percentage input from MBONs and MB2ONs of 0.6% and 11.2%, respectively, suggesting Ipsigoro receives more input from the MB than is typical of a DN.

### Ipsigoro can activate Goro and evoke rolling escape

The structural analysis shown in **Figure 2a-b** suggests that Ipsigoro is a good candidate to mediate descending control of Goro by the MB. To investigate this further we generated a Split-GAL4 line that selectively targets gene expression to Ipsigoro (Ipsigoro-Gal4, **Figure 2d**). Using the line, we first investigated if the connection between Ipsigoro and Goro is functional and, if so, its sign. We expressed the optogenetic activator CsChrimson in Ipsigoro and the fluorescent calcium indicator GCamp6s in Goro in an ex-vivo central nervous system (CNS) preparation. We found that optogenetic activation of Ipsigoro induced a significant increase in the mean Goro ΔF/F, suggestive of an excitatory response (**Figure 2e-f**). This suggests that the synaptic connection between Ipsigoro and Goro is functional and excitatory.

Finally, we wanted to know if activation of Ipsigoro could modulate the rolling escape behaviour. We used the Ipsigoro-Gal4 line to selectively express the temperature sensitive activator dTrpA1 in these neurons and monitored larval behaviour in response to thermogenetic activation of Goro. We found that larvae expressing dTrpA1 in Ipsigoro displayed strong heat-evoked rolling behaviour, significantly more than control animals which lacked the cell-specific enhancers (empty-Split-Gal4 control) and showed very little rolling to heat (**Figure 2g-h**). This suggests that activation of Ipsigoro can promote rolling behaviour.

Consistent with Ipsigoro only driving rolling in a noxious context, optogenetic activation of Ipsigoro was not sufficient to induce the rolling seen to thermogenetic stimulation. However, when combined with noxious stimuli such as heat (used for thermogenetic stimulation) or Basin activation, potentiated rolling was seen in response to optogenetic activation of Ipsigoro (see **Supplementary Figure S1**).

Overall, our results so far suggest that the dominant descending input to the rolling command-like neuron Goro comes from Ipsigoro. This connection was shown to be functional, excitatory and sufficient to evoke a measurable change in rolling behaviour. This suggests that the threshold for activation of Goro by nociceptive cues could be dynamically modulated by synaptic input from the brain. Ipsigoro was shown to receive extensive input from the MB with more than is typical of neurons descending to the VNC and far more than is received by the other descending input to Goro. This suggests that Ipsigoro is specialised to convey learned information about CSs and represents the key mediator of synaptic communication between the MB and Goro. This could allow classically conditioned stimuli to modulate the escape circuitry, allowing CSs to influence escape behaviour as seen in **Figure 1**.

### Encoding of innate context by Ipsigoro

For context-dependent memory-based action selection, animals must integrate learned information with information about co-occurring contextual stimuli. One place this could occur is at the command-like neurons themselves which could receive innate and learned inputs from separate pathways with no prior interaction. Alternatively, integration could occur earlier in the sensorimotor pathway. In the *Drosophila* larva, integration between nociceptive information and learnt information could be carried out by Goro, or by Ipsigoro, or both. To address this, we investigated whether Ipsigoro receives innate input from nociceptive pathways and how this contributes to nociceptive rolling in a naïve animal.

First, we asked if there are structural pathways between nociceptive sensory neurons (MD-IV) and Ipsigoro in the full L1 CNS EM volume, in which the brain has been fully, and the nerve cord partially reconstructed (Ohyama et al., 2015, Winding et al. 2023) and found the shortest pathway consisted of 4 hops (**Figure 3a**): MD-IV neurons synapse onto second order ascending neurons A09e_a3 and A09o_a3 which converge onto a pair of suboesophageal zone (SEZ) neurons, which in turn synapse onto FFN-27, a strong Ipsigoro input which was previously also identified as an input to the vertical lobe DANs that mediate aversive learning (Eschbach et al., 2020).

**Fig. 3.**
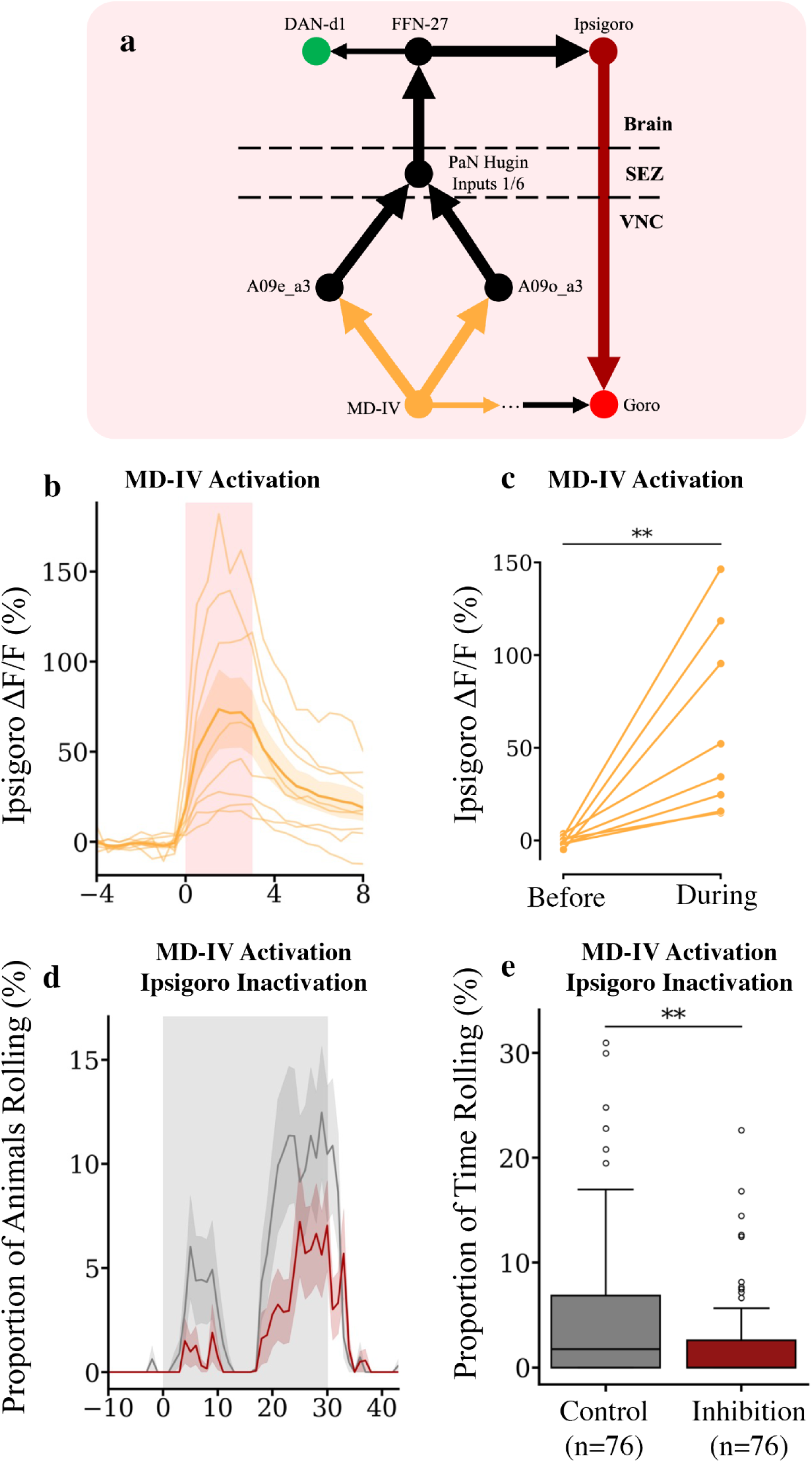
Ipsigoro encodes innate context. a) Structural pathways connect MD-IV neurons to Ipsigoro. The shortest VNC-brain-VNC loop from MD-IV neurons to Goro via Ipsigoro, satisfying the ≥1% input threshold is displayed (bold arrows). Each node represents a left-right pair of homologous neurons except in the case of the 6 A1 MD-IV neurons which are all merged into a single node. Some previously published pathways are displayed with thin arrows including the VNC pathway between MD-IV neurons and Goro and a direct connection between FFN-27 and DAN-d1, a dopaminergic neuron that drives aversive conditioning. This analysis shows that nociceptive input could reach Ipsigoro within 4-hops, suggesting there may be a functional connection. **b-e) Innate pathways to Ipsigoro are functional. b)** Time-series plot of Ipsigoro ΔF/F averaged over three windows of optogenetic MD-IV activation and over three recordings (nine total stimulations). Thin lines show data for each individual animal, the thick line shows the mean of all animals. Data was downsampled to 2Hz to improve visualisation. Red shading indicates the delivery of the red-light stimulus. Error shading shows the mean ± s.e.m. An excitatory response in Ipsigoro is seen during MD-IV stimulation (MD-IV-lexA>lexAop-CsChrimson, Ipsigoro-Gal4>UAS-GCaMP8s). **c)** Mean Ipsigoro ΔF/F is significantly higher during MD-IV stimulation as compared to the 3s prior suggesting the presence of an excitatory pathway. Statistics were conducted using a Wilcoxon signed-rank test; ** p<0.01, N=8. **d)** Time-series plot of the percentage of animals rolling with thermogenetic activation of MD-IV neurons with or without inactivation of Ipsigoro. Data were down-sampled to 1Hz to improve visualisation. Grey shading indicates the delivery of the heat stimulus. Error shading shows the mean rolling ± s.e.m. (MD-IV-lexA>lexAop-dTrpA1, ((Ipsigoro-Gal4 or empty-Split-Gal4)>UAS-shibirets1)). **e)** Boxplot of the percentage of time animals spent rolling during stimulation. Boxes show medians and upper and lower quartiles. Whiskers show upper quartile + 1.5*IQR with outliers plotted as empty circles. There was significantly less heat-evoked rolling in the Ipsigoro inactivation group suggesting this neuron contributes to full expression of nociceptive rolling. Statistics were calculated with a two-sided Mann-Whitney U test; ** p<0.01.

Ascending pathways from MD-IV neurons to MB DANs of over 4 hops have been previously identified and functionally validated, suggesting that the pathways displayed in **Figure 3a** may be functional (Eschbach et al., 2020). To test this, we expressed the fluorescent calcium indicator GCaMP8s in Ipsigoro in an ex-vivo CNS preparation and stimulated CsChrimson expressing MD-IV neurons optogenetically, with red-light. As shown in **Figure 3b-c**, there is a significant increase in the mean Ipsigoro ΔF/F during MD-IV activation. This reveals that the synaptic pathways between MD-IV neurons and Ipsigoro are functional and excitatory.

Next, we wanted to know whether activity in Ipsigoro contributes to the rolling seen in response to a nociceptive stimulus. We activated MD-IV neurons thermogenetically, using dTRPA1 while simultaneously inactivating Ipsigoro using Shibire^ts1^. **Figure 3d-e** shows that inactivation of Ipsigoro significantly reduced rolling to nociceptive stimulus as compared to a control lacking the driver for Ipsigoro, suggesting that descending input from Ipsigoro to the rolling circuitry contributes to nociceptive rolling, likely by virtue of the ascending-descending loop shown in **Figure 3a**.

### Ipsigoro receives functional input from MBONs

For classical conditioning to modulate the selection of a behaviour, it must have a measurable effect on the relevant command-like neurons such that they respond differently when the CS is re-encountered, after learning. The output of the MB is carried by MBONs which are subdivided into approach or avoidance types representing positive and negative value, respectively (Eichler et al., 2017; Eschbach et al., 2021). The neurotransmitter profiles of many MBONs are known including cholinergic, GABAergic and glutamatergic neurons. Cholinergic transmission is generally thought to be excitatory in *Drosophila* although this is not always the case (Bielopolski et al., 2019), while GABAergic transmission is predominantly inhibitory (Lee et al., 2003), whereas glutamatergic could be either. We wanted to analyse in more detail the structural inputs from MBONs onto Ipsigoro and then test them functionally.

3/19 neurons that provide strong, reliable input onto Ipsigoro were MBONs [**Figure 4a** and **Supplementary Table 1**, (Eichler et al., 2017)]. These include the approach-promoting, GABAergic MBON-d1, and the avoidance-promoting, glutamatergic, MBON-k1, and one with untested behavioural effect, due to the lack of a driver line [MBON-b3 (Eschbach et al., 2021)]. 10/19 inputs are MB2ONs [directly downstream of MBONs, (Eschbach et al., 2021)]. Overall, 15/24 MBON pairs were found to form strong inputs to MB2ONs upstream of Ipsigoro corresponding to input from 9/11 of the MB compartments. This suggests that Ipsigoro receives a read-out of the overall population activity of MBONs.

**Figure 4.**
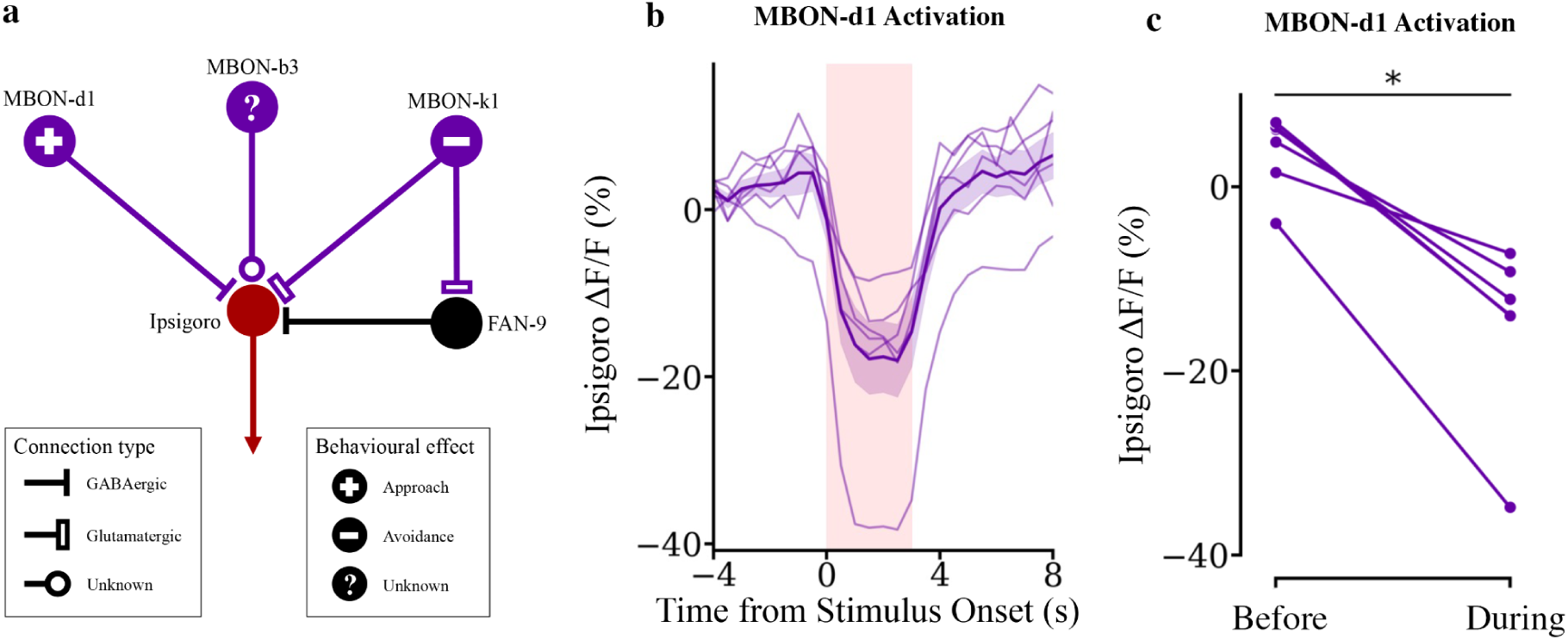
Ipsigoro receives functional input from the Mushroom Body. **a)** Ipsigoro receives strong direct synaptic input from 3 MBONs. Neurotransmitter profiles and behavioural effects are from previous publications (Eichler et al., 2017; Eschbach et al., 2020). GABAergic and Glutamatergic inputs are expected to be inhibitory. MBON-d1 is approach-promoting and GABAergic, it is therefore predicted to inhibit Ipsigoro. MBON-k1 is avoidance-promoting and glutamatergic. MBON-b3 has an unknown behavioural effect and neurotransmitter profile. For one two-step pathway from these MBONs, the neurotransmitter profile of the intermediate neuron is known (FAN-9, GABAergic) suggesting that this pathway from MBON-k1 is disinhibitory. This suggests the functional connectivity between MBON-k1 and Ipsigoro may be complex with the direct pathway predicted to be inhibitory and the indirect pathway disinhibitory. **b)** Time-series plot of Ipsigoro ΔF/F averaged over three windows of optogenetic MBON-d1 activation and over three recordings (nine total stimulations). Thin lines show data for each individual animal, the thick line shows the mean of all animals. Data was downsampled to 2Hz to improve visualisation. Red shading indicates the delivery of the red-light stimulus. Error shading shows the mean ± s.e.m. An inhibitory response in Ipsigoro is seen during MBON-d1 stimulation (MBON-d1-lexA>lexAop-CsChrimson, Ipsigoro-Gal4>UAS-GCaMP8s). **c)** Mean Ipsigoro ΔF/F is significantly lower during MBON-d1 stimulation as compared to the 3s prior suggesting the presence of an excitatory pathway. Statistics were conducted using a Wilcoxon signed-rank test; * p<0.05, N=6.

Next, we wanted to functionally verify the connections between the MB and Ipsigoro. A previously identified MBON-d1-lexA line allowed the strongest MBON input to be functionally tested (Meissner et al. 2024). Using this line, we expressed CsChrimson in MBON-d1 and the fluorescent calcium indicator GCaMP8s in Ipsigoro. Optogenetic activation of MBON-d1 with red-light evoked inhibitory responses in Ipsigoro (**Figure 4b-c)**. This fits with the predicted sign of the direct synaptic connection shown in **Figure 4a** and shows that the MB output can modulate Ipsigoro as suggested by the connectivity.

### Goro command neurons receive functional input from MBONs, which bi-directionally toggle its activity

Finally, we also wanted to know if the inputs from MBONs could drive measurable responses in Goro. Using a Goro-lexA line (Ohyama et al., 2015) along with split-Gal4 lines for MBON-d1 and MBON-k1 (Eschbach et al. 2021; Saumweber et al. 2018), we were able to test connections from both of these neurons. **Figure 5c-d** shows that optogenetic activation of CsChrimson expressing MBON-d1 and MBON-k1 was able to evoke inhibition and excitation of Goro, respectively. Thus, the two MBONs that encode opposite learnt valence toggle Goro activity in opposite ways: the approach-promoting MBON-d1 that encodes positive valence, inhibits Goro (and therefore reduces the likelihood of roll-escape); the avoidance promoting MBON-k1 that encodes negative value activates Goro (and therefore increases the likelihood of roll-escape).

**Figure 5.**
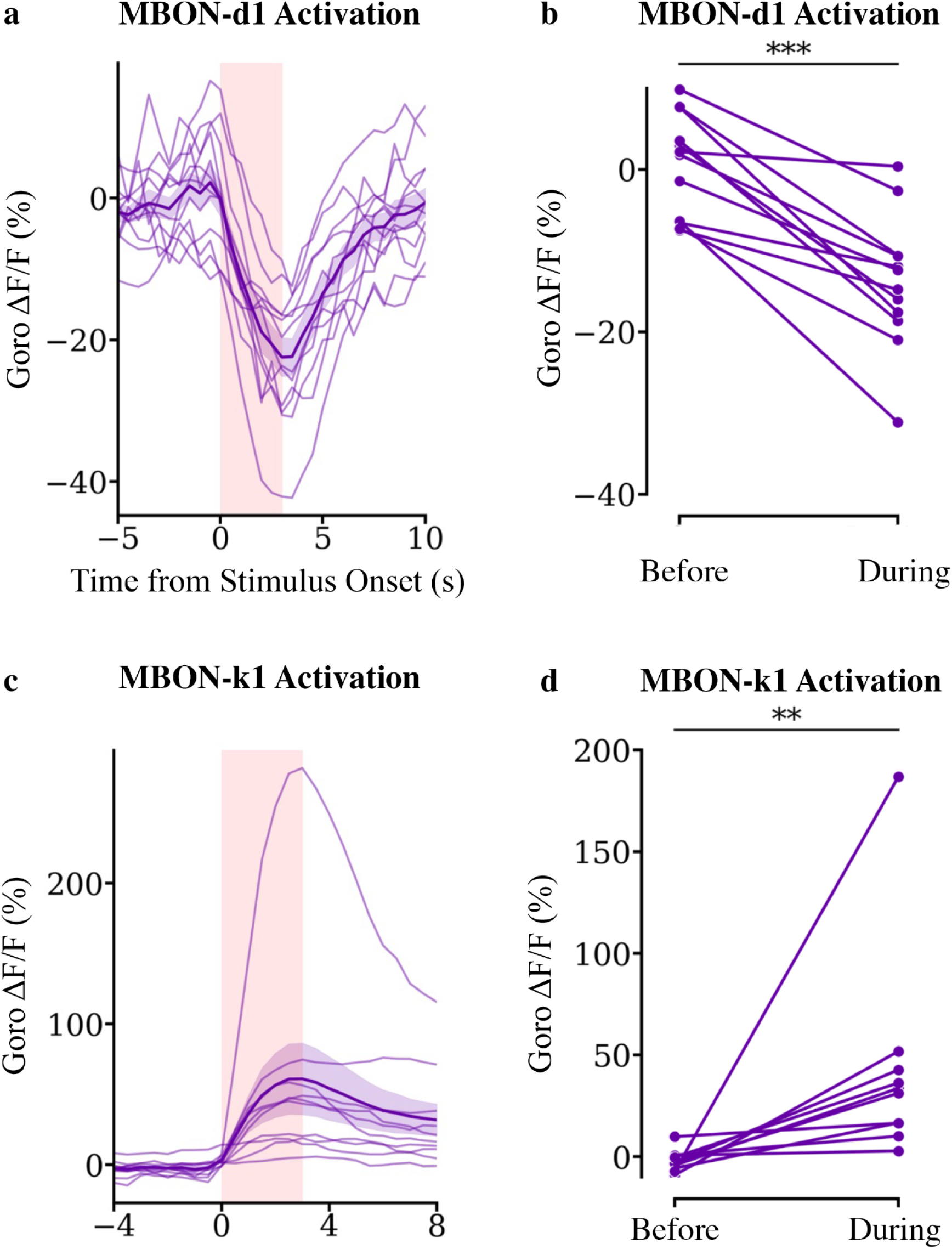
Activation of MBON-d1 (Approach) and MBON-k1 (Avoidance) evokes opposite responses in Goro. **a)** Time-series plot of Goro ΔF/F averaged over three windows of optogenetic MBON-d1 activation and over three recordings (nine total stimulations). Thin lines show data for each individual animal, the thick line shows the mean of all animals. Data was downsampled to 2Hz to improve visualisation. Red shading indicates the delivery of the red-light stimulus. Error shading shows the mean ± s.e.m. An inhibitory response in Goro is seen during MBON-d1 stimulation. (MBON-d1-Gal4>UAS-CsChrimson, Goro-lexA>lexAop-GCaMP6s). **b)** Mean Goro ΔF/F is significantly lower during MBON-d1 stimulation as compared to the 3s prior suggesting the presence of an inhibitory pathway. Statistics were conducted using a two-sided Wilcoxon signed-rank test; *** p<0.001, N=12. **c)** Time-series plot of Goro ΔF/F averaged over three windows of optogenetic MBON-k1 activation and over three recordings (nine total stimulations). Thin lines show data for each individual animal, the thick line shows the mean of all animals. Data was downsampled to 2Hz to improve visualisation. Red shading indicates the delivery of the red-light stimulus. Error shading shows the mean ± s.e.m. An excitatory response in Goro is seen during MBON-k1 stimulation. (MBON-k1-Gal4>UAS-CsChrimson, Goro-lexA>lexAop-GCaMP6s). **d)** Mean Goro ΔF/F is significantly higher during MBON-k1 stimulation as compared to the 3s prior suggesting the presence of an excitatory pathway. Statistics were conducted using a two-sided Wilcoxon signed-rank test; ** p<0.01, N=10.

The inhibition of Ipsigoro and Goro by MBON-d1, demonstrated here, presents a possible circuit mechanism for aversive learning to modulate its activity (**Figure 6a-d**). MBON-d1 innervates the lateral appendix compartment of the MB which receives input from DAN-d1. DAN-d1 receives functional input from nociceptive pathways and supports aversive odour memory (Eschbach et al., 2020). Interestingly it is one of two DANs (-d1 and-g1) which have been shown to be functionally connected to the Basin neurons themselves (Eschbach et al., 2020). Previous work in the adult and larval *Drosophila* shows that aversive conditioning depressed conditioned odour drive to approach promoting MBONs (Hige et al., 2015; Séjourné et al., 2011; Eschbach et al. 2021). Pairing of odours with Basin activation would therefore depress CS-evoked responses in MBON-d1, resulting in less inhibition of Ipsigoro when the odour is re-encountered (**Figure 6b-d**). **A**pproach promoting MBON-d1 and the avoidance promoting MBON-k1 drive inhibitory and excitatory responses in Goro respectively (**Figures 5a-d and 6a-c**). Given that Ipsigoro is the dominant descending input to Goro and is the descending input that receives by far the strongest inputs from MB neurons (see **Figure 2a-b**), these effects are likely through modulation of this neuron. In a naïve animal, excitatory and inhibitory MB influences over Goro would be predicted to balance each other out (**Figure 6a-c**). However, after learning, the depression of MBON-d1 responses to odour would result in a net excitation and therefore a greater probability of rolling in the presence of the CS (**Figure 6b-d**). In the absence of nociceptive drive to Ipsigoro and Goro, the excitatory drive to Goro from the MB would not be sufficient to evoke rolling (**Figure 6b**). However, the presence of nociceptive context, nociceptive input to Ipsigoro and Goro would be integrated with the excitatory input from the MB to increase the likelihood of rolling (**Figure 6d**).

**Figure 6.**
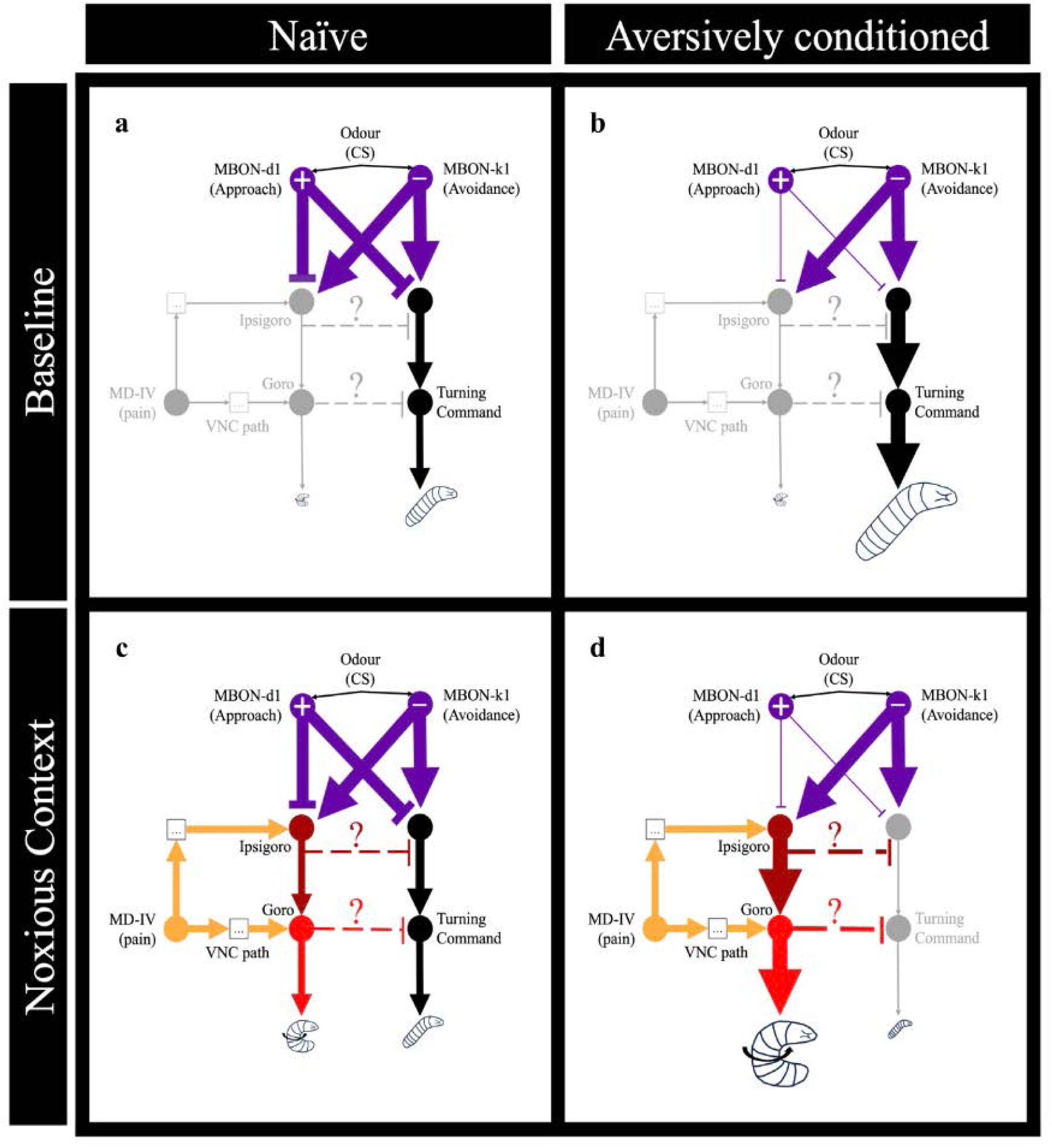
Schematic showing how the identified circuit could function during context-dependent memory-based action selection. Nodes in the circuit represent homologous left-right pairs of neurons (except in the case of the larger group of MD-IV neurons). “VNC path” refers to the previously published VNC based pathway between MD-IV neurons and Goro (Ohyama et al., 2015). **a)** At baseline, a naïve animal navigates its environment by probabilistic transitions to turning behaviours under the control of an unidentified descending pathway (black). Odour (conditioned stimulus, CS) activates MBONs but the approach-and avoidance-promoting systems provide a balanced input to the turning command system. Activity in the VNC path and in Ipsigoro is too weak too activate Goro and evoke rolling. **b)** After aversive conditioning, input from the approach promoting MBON is reduced, disinhibiting the turning command such that there is more turning in the presence of the odour CS. Activity in the VNC path and in Ipsigoro remain low in the absence of nociceptive input and therefore disinhibition of Ipsigoro does not evoke a measurable change in rolling. **c)** In the noxious context, the VNC pathway and ascending inputs to Ipsigoro become active, setting a baseline of activity in Ipsigoro and Goro with an associated increase in rolling. Rolling is expected to suppress turning behaviour through an unidentified inhibitory pathway. **d)** After aversive conditioning, the baseline of activity set in Ipsigoro in the noxious context can be modulated. Depression of input from the approach promoting MBON disinhibits Ipsigoro driving an increase in rolling, again at the expense of turning.

## Discussion

Even in the most basic centralised nervous systems, there is a clear subdivision between sensory input and motor output, allowing classical conditioning to occur. In organisms like nematodes and simple molluscs, interneurons act to integrate sensory signals and organise motor neurons to generate discrete behaviours (Hochner & Glanzman, 2016; White et al., 1997). Action selection is underpinned by command neurons which typically receive synapses from sensory neurons and project directly to motor neurons (Kupfermann & Weiss, 1978). Even in this simple arrangement, classical conditioning is possible and involves key principles that are adapted in higher organisms. For example, in the terrestrial snail *helix*, pairing of a food CS with electric shock results in the formation of a conditioned withdrawal response, with an increased responsiveness of withdrawal command neurons to the paired food (Balaban et al., 1987). Neuromodulatory teaching signals play a key role, with stimulation of a single identified nociceptive serotonergic cell being sufficient to induce these changes, likely through plasticity in the sensory inputs to the command neuron (Balaban et al., 2001; Zakharov et al., 1995).

In higher organisms such as vertebrates, there is an increase in complexity with an increased number of command-like neurons which are often separated from their sensory inputs by functionally specialised networks of interneurons (Jing, 2009). Rather than modulating direct links between sensory neurons and command neurons, neuromodulators such as dopamine guide classical conditioning in specialised learning centres which first assign CSs with a value which can then be used flexibly to inform many context-dependent behaviours (Schultz, 2005). The outputs of learning centres such as the striatum and the cerebellum are carried by GABAergic projection neurons, with control of behaviour thought to involve shifts in the balance of inhibition and disinhibition (Marr, 1969; Mink, 1996). For example, subpopulations of striatal inhibitory projection neurons contributing to “direct” and “indirect” output pathways have been found to drive excitatory and inhibitory responses in glutamatergic neurons of the mouse Mesolimbic Locomotor Region (MLR), a key command centre for locomotor behaviour (Roseberry et al., 2016). Centres such as this reach motor circuits in the nerve cord by virtue of descending neurons from the brain stem (Grillner & El Manira 2019; Juvin et al. 2016). In some cases, descending neurons themselves are command-like, projecting directly to motor neurons (Korn & Faber 2005). In other cases, command systems are found in the spinal cord allowing behaviour patterns to be expressed even following spinalization (Rossignol et al., 2002). Only learned behaviours relevant to the current context should be expressed, requiring learned pathways to be integrated with innate pathways prior to action selection. The circuits that perform this integration, allowing learning centres to influence command-like neurons in a context-dependent manner are not well understood.

Arthropod nervous systems offer a useful intermediate in complexity between simpler invertebrates and complex vertebrates. They also have specialised learning centres and extensive networks of descending neurons guiding behaviour but their circuits are small enough to be studied at the level of individual neurons and synapses (Cande et al., 2018; Winding et al., 2024). Here, we demonstrate *Drosophila* larval escape can be modulated by associative learning and identify the circuitry that mediates context-dependent memory-based action selection (**Figure 6)**.

This circuit has several key properties that can be compared to vertebrate systems. Firstly, aversive conditioning can modulate selection of an escape behaviour that is not seen outside of the noxious context. This suggests the behavioural consequences of negatively valued stimuli are flexible and context-dependent as in vertebrates (McDannald, 2023). Secondly, there is a multipath architecture underpinning nociceptive activation of the escape command neurons. This includes the previously published short VNC pathway via Basin neurons and a longer pathway via an excitatory descending neuron (Ohyama et al., 2015). This is reminiscent of vertebrate escape behaviour where the short pathways seen in simple nervous systems are retained to allow rapid triggering of the behaviour in the presence of strong aversive stimulation (J. D. Robertson et al., 1963). Thirdly, the command-like neurons receive functional inhibitory and excitatory pathways from positive and negative value-coding learning centre outputs (MBONs). Both these MBONs have inhibitory neurotransmitter profiles suggesting that learning modulates the balance of inhibitory and disinhibitory projections as proposed in the vertebrate striatum and cerebellum (Marr, 1969; Mink, 1996). Fourthly, these pathways involve the same descending neuron that receives ascending nociceptive information, allowing integration of contextual and learned information. This could allow aversive memories to only evoke the behaviour in the relevant innate context. Finally, the learned inputs and innate inputs also synapse onto the same DAN (DAN-d1) expected to evoke the memory. Input from innate pathways may suggest that this DAN and associated learning centre compartment could be specialised for escape learning, receiving escape-relevant teaching signals. Input from learned pathways suggests that learned information is both used to inform escape action selection and to inform future learning that is relevant to control of the escape behaviour. This dual function of learned value signals is commonly seen in solutions to RL problems (Sutton & Barto, 1998).

All together the results presented in this study reveal a synaptic level circuit underpinning context-dependent memory-based action selection in the *Drosophila* larva. These results extend the current understanding of how learned value outputs of learning centres like the MB are used to guide action selection. Results highlight the importance of descending inputs to command neurons in the integration of opposing positive and negative value inputs from the MB with ascending information about innate context.

## Acknowledgements

The authors thank Nan Hu and Oxana Elliott at the Department of Zoology, University of Cambridge for fly crosses; AT for very helpful comments on the manuscript; MRC Laboratory of Molecular Biology Core funding (MZ, BMWJ, NGC, SNH), Wellcome Trust grant 205050/Z/16/Z (MZ), ERC grant ERC-2018-COG: 819650 (MZ) for funding.

## Materials and Methods

### Fly Stocks

All animals used were third instar wandering stage larvae of the *Drosophila melanogaster* species. Larvae were reared on fly food with the following composition: molasses 41.7ml, yeast 21.0g, corn meal 87.1g, agar 7.0g, 10% Nipagin 10.4ml, Propionic acid 5.2ml in 1L of water. For experiments involving optogenetic activators, food was supplemented with 5ml of a 100mM all-trans-retinal stock per 1L of medium. All animals were reared in the dark. For imaging experiments, animals were reared at 25°C. For behavioural experiments (where thermogenetics was generally used), animals were reared at 18°C to avoid neuronal manipulations during larval development.

The GAL4/UAS and LexA/LexAop binary expression systems were used, either alone or in combination, to target neurons (Brand & Perrimon, 1993; Lai & Lee, 2006). The specific expression systems used for experiments are highlighted in the corresponding figure legends. Optogenetic activation was achieved using targeted CsChrimson expression (Klapoetke et al., 2014). Thermogenetic activation and inactivation were achieved using targeted expression of dTrpA1 and Shibire^ts1^ respectively (Kitamoto, 2001; Rosenzweig et al., 2005). The Or42b driver was used to target expression in a subset of olfactory neurons for learning experiments (Fishilevich & Vosshall, 2005). R72F11 and *ppk* drivers were used for expression in Basin and MD-IV neurons respectively (Ainsley et al., 2003; Ohyama et al., 2015). For expression in Ipsigoro the SS03731 driver was used (Meissner et al., 2024). For expression in MBON-d1, SS01705 and VT007911 drivers were used (Eschbach et al., 2021; Meissner et al., 2024; Tirian & Dickson, 2017). For expression in MBON-k1, the SS01962 driver was used (Saumweber et al., 2018). A complete list of fly stocks used is displayed in **Supplementary Table 2** with the listed abbreviated names used to refer to lines in the remainder of this section.

### Functional connectivity experiments

All functional connectivity experiments were conducted in a dissected CNS preparation adhered to a poly-lysine coated coverslip by the VNC with the brain lobes pointing upwards. Samples were mounted in the lid of a 35×10mm petri dish and immersed in Baines external solution (Marley & Baines, 2011) with the following composition: 135 mM NaCl, 5 mM KCl, 4 mM MgCl_2_, 2 mM CaCl_2_, 5 mM TES and 8.7mM Glucose. The pH was adjusted to 7.15 using NaOH with an osmolarity of 310-320 mOsm. Postsynaptic neuronal responses were recorded by imaging fluctuations in the fluorescence of the green calcium indicators GCaMP6s (recordings in Goro) or GCaMP8s (recordings in Ipsigoro) (Chen et al., 2013; Zhang et al., 2023). Imaging was performed using a 3i VIVO multiphoton upright microscope (Intelligent Imaging Innovations) with a 2-photon imaging laser tuned to 920nm (Insight DS+ Dual, Spectra-Physics) and an Apo LWD 25x/1.10W objective (Nikon). Fast imaging was achieved using resonant scanning (Vector RS+). Data was collected on a single Z-plane that captured the dendrites of the postsynaptic neuron. In the case of Goro, dendrites could not be reliably separated from axons so recorded fluorescence likely originated from both these structures. Data was captured at 39.7 frames/s (recordings in Goro) or 9.9 frames/s (recordings in Ipsigoro). The imaging power measured at the objective was between 10 and 20 mW with the precise value selected prior to use of each new genotype to ensure a clear image.

Optogenetic stimulation of presynaptic neurons was delivered using a 640nm laser (LaserStack 3iL33) coupled to a 1-photon Phasor to restrict stimulation to a specified holographic pattern. An oval photo-stimulation region of dimensions 140µm x 75µm was centred over the brain hemispheres (stimulation of Ipsigoro or MBONs) or the anterior segments of the VNC (stimulation of MD-IV neurons) depending on the location of the presynaptic neurons. For all experiments the same stimulation protocol was used. Each recording lasted a total of 120s. After an initial rest of 30s, samples underwent 3 stimulation trials delivered at 30s, 60s and 90s. At the start of each trial 3s of red-light stimulation was delivered followed by 27s of rest. Three recordings were taken for each sample. The stimulation power measured at the objective was 20 µW.

For the recordings in Ipsigoro, females from the Ipsigoro-Gal4>UAS-GCaMP,lexAop-CsChrimson line were crossed with males from the MD-IV-lexA (MD-IV stimulation) or MBON-d1-lexA (MBON-d1 stimulation) lines. For recordings in Goro, females from the Goro-lexA>lexAop-GCaMP,UAS-CsChrimson line were crossed with males from the Ipsigoro-Gal4 (Ipsigoro stimulation), MBON-d1-Gal4 (MBON-d1 stimulation) or MBON-k1-Gal4 (MBON-k1 stimulation) lines. In the case of the MBON-d1-lexA line, expression in MBON-d1 was stochastic so each brain was screened for expression prior to being used for experiments. Prior to the use of each photo-stimulation region location and postsynaptic neuron combination, an effector control was conducted where w^1118^ males were crossed to females from the Goro-lexA>lexAop-GCaMP,UAS-CsChrimson line (recordings from Goro) or the Ipsigoro-Gal4>UAS-GCaMP,lexAop-CsChrimson line (recordings from Ipsigoro) to ensure no signal was observed in the absence of targeted CsChrimson expression. Eggs were collected on retinal food and incubated in the dark at 25°C for 4 days prior to recording.

### Analysis of calcium imaging data

Imaging data was processed using Fiji (Schindelin et al., 2012). A region of interest (ROI) was manually defined to include the postsynaptic GCaMP-expressing dendrites. The average pixel value inside the ROI was measured. The mean pixel value in the remainder of the image was subtracted from this value to give a background-subtracted fluorescence value for each time point. These data were analysed using custom code written in Python3 particularly using the Pandas package (McKinney, 2010; Van Rossum & Drake, 2009). Fluorescence values were averaged across stimulation trials and across recordings for each sample to give a single fluorescence time series representing the mean fluorescence (F) across a total of 9 stimulation trials. Mean fluorescence in the 15s preceding stimulation was taken as a measure of baseline fluorescence (F_0_) for the sample. Changes in fluorescence were quantified as (F-F_0_)/F_0_ to give a ΔF/F value at each timepoint. Imaging data was plotted using the seaborn and matplotlib Python packages (Hunter, 2007; Waskom, 2021).

### Behavioural experiments

The apparatus used for behavioural experiments is described in detail in our previous publication (Croteau-Chonka et al., 2022). In short, larvae are placed on a 23cm x 23cm 3% agarose plate within a dark enclosure. The plate is illuminated below by an 850nm LED backlight (#SOBL-300×300–850, Smart Vision Lights) such that larval outlines can be captured by an overhead camera at 20 frames/s (#TEL-G3-CM10-M5105, Teledyne DALSA). For optogenetic stimulation, uniform 640nm LED light is delivered to the plate using two digital micromirror devices (#296-DLP4710EVM-LC-ND, Texas Instruments). Visible wavelengths are prevented from reaching the camera by an 800nm long-pass filter (#LP800-40.5, Midwest Optical Systems). Thermal stimulation is delivered by a 1470nm infrared laser (#LRD-1470-PFI-15000–05, Laserglow Technologies) directed through a two-axis scanning galvanometer (#GVSM002, Thorlabs). The high scanning velocity of the galvanometer allows rapid cycling between larvae to allow fast simultaneous heating. A thermal camera (Teledyne FLIR AX8) is installed for online adjustment of laser power such that a desired temperature can be maintained.

Camera images were processed in parallel by a host computer (#T7920, running Windows 10, Dell Technologies Inc) and an associated field programmable gate array (FPGA) device (#PCIe-1473R-LX110, National Instruments). Software controlling stimulation and image processing used LabView and C++ (Bitter et al., 2006; Stroustrup, 1995) and can be found at https://github.com/ZlaticLab/multi-larva-tracker-scripts-public (Clayton et al., 2022). Larval outlines and spines were extracted with spines consisting of 11 points along the central body axis running from head to tail (Croteau-Chonka et al., 2022; Swierczek et al., 2011). Additional corrections to head/tail orientation and removal of distorted traces falling outside of reasonable area bounds were conducted using custom Python3 software (Van Rossum & Drake, 2009) written by Peter Hague in our laboratory. Data in **Supplementary Figure S1b** and **S1c** were generated using a higher throughput optogenetic stimulation apparatus (∼30 animals per experimental run). It has a similar arrangement, but stimulation light is of 625nm and is delivered by an array of LEDs from below. This arrangement has been described previously (Jovanic et al. 2016; Ohyama et al. 2015).

Prior to experiments, eggs were collected and incubated in the dark at 18°C either for 7 days on retinal food (learning) or 8 days on non-retinal (Ipsigoro activation and inactivation) or retinal (**Supplementary Figure S1**) food. Before each experimental run, 8 larvae were removed from food, washed with water and transferred to the agarose plate with a small paintbrush. At the start of the run, animals were spread out to fill a circle of diameter ∼10cm in the centre of the plate to reduce the chance of collision events.

For learning experiments, males from the Basin-Gal4>UAS-dTrpA1 line were crossed to females of the Or42b-lexA>lexAop-CsChrimson line to allow optogenetic activation of Or42b neurons (CS) and thermogenetic activation of Basin neurons (US). The full training and testing protocol is displayed in **Figure 1a**. After a pre-test period of 30s, two groups of animals received 24 training trials on an agarose plate. All stimulations lasted 20s. For training, animals were divided into forward-and backward-paired groups. For photo-stimulation, a red-light intensity of 550 µW/cm^2^ measured at the agar plate was used. For thermal stimulation, a temperature of 27.5°C was used to activate Basin neurons, producing high levels of rolling as seen previously (Ohyama et al., 2015). In the forward-paired group, thermal stimulation followed olfactory stimulation with a 1s gap in between. In the backward-paired group, optogenetic stimulation followed punishment with a 5s gap. After training, animals were transferred to a testing agarose plate and after a 30s pre-test period, photo-and thermal stimulation were delivered simultaneously. During the test a red-light intensity of 550 µW/cm^2^ was again used but animals were instead heated to 25.1°C. This lower temperature was chosen to produce a more moderate level of rolling that could be modulated by learning. The threshold for dTrpA1-evoked spiking in *Drosophila* neurons has been previously shown to be around 25°C (Pulver et al., 2009). Three test stimulation trials were delivered but all analysis was restricted to the first stimulation due to the possibility of additional conditioning occurring across trials.

For thermogenetic activation of Ipsigoro, females from the UAS-dTrpA1 line were crossed with males from the Ipsigoro-Gal4 (Ipsigoro activation) or empty-Split-Gal4 (control) lines. After a rest period of 30s, larvae were heated to 27.5°C for 20s on a moist agarose plate followed by a rest of 60s. For thermogenetic inactivation of Ipsigoro during MD-IV stimulation, females from the MD-IV-lexA>lexAop-dTrpA1,UAS-Shibire^ts1^ line were crossed to males from the Ipsigoro-Gal4 (Ipsigoro inactivation) or empty-Split-Gal4 (control) lines. After a rest period of 30s, larvae were heated to 30.1°C for 30s on a moist agarose plate followed by a rest of 60s. Three stimulation trials were delivered for both activation and inactivation experiments but all analysis was restricted to the first stimulation due to habituation of the behaviour.

For the experiments analysed in **Supplementary Figure S1a**, females from the UAS-CsChrimson line were crossed with males from the Ipsigoro-Gal4 (Activation Only and Activation + Heat groups) or empty-Split-Gal4 (Heat + Light group) lines. For thermal stimulation, animals were heated to 27.5°C. For optogenetic activation/light stimulation, animals were exposed to red-light with an intensity of 800 µW/cm^2^ measured at the agar plate. After a rest period of 30s, larvae were stimulated for 20s on a moist agarose plate followed by a rest of 90s. For the data in **Supplementary Figure S1b** and **S1c,** females from the Basin-Gal4>UAS-CsChrimson lines were crossed with males from the Ipsigoro-Gal4 (Ipsigoro activation) or empty-Split-Gal4 (control) lines. After a rest period of 30s, larvae were exposed to red-light with an intensity of 75 µW/cm^2^ for 30s an agarose plate followed by a rest of 30s. This low intensity was selected to evoke moderate levels of rolling in the Basin stimulation-only control. Again, multiple stimulation trials were delivered but all analysis was restricted to the first stimulation window.

### Analysis of behavioural data

Rolling data were generated using the LarvaTagger software (Laurent et al., 2024). LarvaTagger allows both manual and automatic tagging of larval behaviours. Only animals present for 95% of the analysed time window were included in analysis. For the learning experiments, roll events were manually tagged using the associated graphical user interface (GUI). For other experiments, an automatic tagger was trained on a dataset consisting of ∼70% of the data from the learning experiments. An additional data file was also manually tagged and included in the training set which consisted of 8 larvae undergoing two 20s windows of strong Basin stimulation. LarvaTagger uses a self-supervised autoencoder, pre-trained on large data repositories accumulated at Janelia Research Campus (Jovanic et al., 2016; Masson et al., 2020; Vogelstein et al., 2014) to project short behavioural sequences into a low-dimensional latent space. These latent representations are then used as input to train a new tagger. This arrangement allows for “transfer learning” such that relatively little new annotated data is required for good performance. Best performance was found when training a classifier to differentiate the C-bend and sliding portions of the roll which were then combined for subsequent analysis. An additional threshold was applied on speed perpendicular to the body axis to reduce false positives. This variable has been termed “crabspeed” in previous publications and is strongly correlated with rolling behaviour (Ohyama et al., 2013). Crabspeed and area for each larval outline was computed using custom Python3 software (Van Rossum & Drake, 2009) written by Andrey Stoychev in our laboratory designed to replicate the crabspeed calculation implemented in the MWT software package (Ohyama et al., 2013; Swierczek et al., 2011). Detected roll events were removed if the 95^th^ percentile event crabspeed (mm/s) failed to exceed 2/3 of the median area of the larval trace (mm^2^). The remaining ∼ 30% of learning data was used to quantify the performance of the algorithm. When computing precision, detected events lying within 0.5s of a true event were still considered to be true positives. When computing recall, detected events had to overlap with the true event to be considered true positives. Using these criteria, the classifier performed with a precision of 0.80 and a recall of 0.71. Data in **Supplementary Figure S1b** and **S1c** from the alternative optogenetic apparatus were analysed using an existing alternative tagger previously trained on data from a similar apparatus (Masson et al. 2020). Behavioural data was processed and plotted largely using the pandas, seaborn and matplotlib Python packages (Hunter 2007; McKinney 2010; Waskom 2021).

### Statistical analysis

Statistical analysis was performed using the Scipy package in Python3 (Van Rossum & Drake, 2009; Virtanen et al., 2020). For functional imaging experiments, the mean ΔF/F during the 3s stimulation period was compared to 3s prior to stimulation using a two-sided Wilcoxon signed-rank test. For behavioural assays, the percentage of time animals spent rolling during stimulation was compared between control and experimental animals using a two-sided Mann-Whitney U test. In the case of the learning, rolling to Basin stimulation occurred almost entirely within a 5s window starting at 4s after the onset of heating. As a result, analysis was restricted to this window. In all other behavioural experiments roll events were spread across the window so the entire stimulation window was considered. For **Supplementary Figure S1a**, where more than one pair of conditions were compared, statistics were calculated using a Kruskal Wallis H test followed by pairwise comparisons with a post-hoc Dunn test with Bonferroni correction. This analysis was performed using the pingouin and scikit_posthocs packages in Python3 (Terpilovskii 2019; Vallat 2018; Van Rossum & Drake 2009). In all figures *, **, *** and **** represent p-values of <0.05, <0.01, <0.001 and <0.0001 respectively with n.s. used to indicate no significant difference.

### Connectomic analysis

All structural circuit analysis used the same L1 larval CNS connectome (available at https://l1em.catmaid.virtualflybrain.org/?pid=1) of which the brain is now fully reconstructed (Winding et al., 2024). Connectivity was analysed using the online software CATMAID (Saalfeld et al., 2009) and custom code written in Python3 (Van Rossum & Drake, 2009). Some weak connections (particularly those of <3 synapses) could be non-functional, transient or erroneous so identified circuits were restricted to synaptic pathways that were bilateral with each connection contributing an average of ≥1% of input to the postsynaptic neuron when averaged between the two sides (Schneider-Mizell et al., 2016). Similar thresholds have been used in previous publications for axo-dendritic connections but here all connection types were considered together (Winding et al., 2024). An exception is **Figure 2b** and **2c** where all synapses were considered to give a percentage of total input from the MB.

## Notes

### Competing Interest Statement

The authors have declared no competing interest.

