## Supplementary Information for "Descending neurons integrate learnt information from mushroom body with context to promote escape behaviour"

#### Supplementary Figure 1

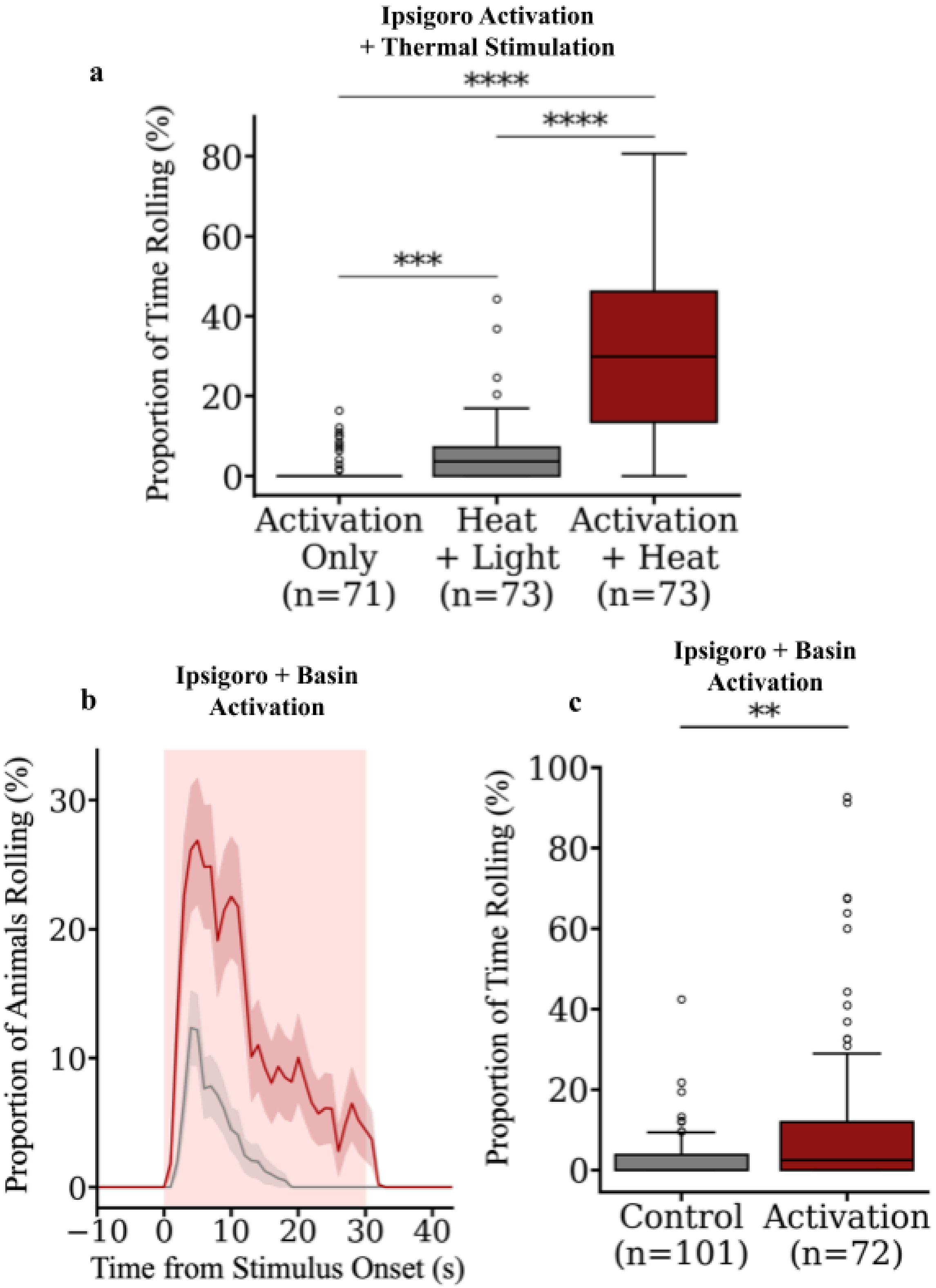

**Supplementary Figure S1. Photostimulation of Ipsigoro potentiates nocifensive rolling.**

**a)** Boxplot of the percentage of time animals spent rolling during a 20s stimulation window. Boxes show medians and upper and lower quartiles. Whiskers show the range of data and, in the case of outliers, the upper whisker shows the upper quartile + 1.5*IQR, with outliers plotted as empty circles. Statistics were calculated using a Kruskal Wallis H test ($p<0.0001$) followed by pairwise comparisons with a post-hoc Dunn test with Bonferroni correction; *** $p<0.001$, **** $p<0.0001$. Optogenetic activation of Ipsigoro alone (Activation group) evokes very little rolling. Simultaneous heat and light delivery to an empty driver control (Heat + Light group) causes a mild increase in rolling. Optogenetic activation of Ipsigoro with simultaneous delivery of a heat stimulus (Activation + Heat group) evokes significantly more rolling than is seen in both the other groups suggesting Ipsigoro specifically drives rolling in a noxious context. ((Ipsigoro-Gal4 or empty-Split-Gal4)>UAS-CsChrimson). **b)** Time-series plot of the percentage of animals rolling in response to optogenetic activation of Basin neurons with or without optogenetic activation of Ipsigoro. Data were down-sampled to 1Hz to improve visualisation. Red shading indicates the delivery of the light stimulus. Error shading shows the mean rolling ± s.e.m. ((Ipsigoro-Gal4 or empty-Split-Gal4), Basin-Gal4>UAS-CsChrimson). **c)** Boxplot of the percentage of time animals in **b** spent rolling during the 30s stimulation window. Boxes show medians and upper quartiles. Whiskers shows the upper quartile + 1.5*IQR, with outliers plotted as empty circles. There was significantly more rolling in the Ipsigoro activation group suggesting this neuron facilitates rolling escape in response to Basin stimulation. Statistics were calculated with a two-sided Mann-Whitney U test; ** $p<0.01$.

### Supplementary Tables

#### Supplementary Table 1

| Name | Cell Type | Ipsilateral | Contralateral | Skeleton IDs | Reference |
|---|---|---|---|---|---|
| MBON-b3 | MBON | | 1.9% | 16883909, 4234009 | (Eichler et al., 2017) |
| MBON-d1 | MBON | 3.4% | 2.4% | 7055857, 4241237 | (Eichler et al., 2017) |
| MBON-k1 | MBON | | 1.2% | 16846805, 18028397 | (Eichler et al., 2017) |
| FAN-9* | FBN | 1.4% | 2.1% | 14260575, 16987409 | (Eschbach et al., 2020) |
| FBN-23* | FBN | | 3.5% | 7026287, 7980369 | (Eschbach et al., 2020) |
| FBN-28* | FBN | | 1.7% | 10612094, 12122924 | (Eschbach et al., 2020) |
| CN-39 | CN | 3.6% | | 12587013, 9015568 | (Eschbach et al., 2021) |
| CN-41 | CN | 1.4% | | 17276867, 6611100 | (Eschbach et al., 2021) |
| MB2ON-35 | MB2ON | 1.4% | | 16445338, 16350988 | (Eschbach et al., 2021) |
| MB2ON-57 | MB2ON | | 1.4% | 6298455, 6193472 | (Eschbach et al., 2021) |
| MB2ON-59 | MB2ON | | 3.8% | 7700444, 16910799 | (Eschbach et al., 2021) |
| MB2ON-74 | MB2ON | | 1.6% | 14539753, 12962772 | (Eschbach et al., 2021) |
| MB2ON-232 | MB2ON | 1.2% | | 14329745, 19786464 | (Eschbach et al., 2021) |
| FB2IN-5 | FB2N | | 1.6% | 17177005, 8019816 | (Eschbach et al., 2020) |
| FFN-1* | FFN | 1.4% | | 9048462, 10612630 | (Eschbach et al., 2020) |
| FFN-23 | FFN | | 2.4% | 16385470, 18098174 | (Eschbach et al., 2020) |
| FFN-27* | FFN | | 2.6% | 14493841, 11361875 | (Eschbach et al., 2020) |
| FB-36 | N/A | | 1.6% | 11871574, 11102156 | (Winding et al. 2024) |
| MB2IN-135 | N/A | 1.2% | | 10323628, 8370111 | (Winding et al. 2024) |

* Previously published as providing input to DAN-d1

**Supplementary Table 1. Ipsigoro receives strong inputs from 19 brain neurons.**

Ipsigoro receives strong pathways (bilateral, mean input percentage ≥1%) from 19 brain neurons. Of these 3 are MBONs, 10 are MB2ONs (3 FBN, 2 CN, 5 other), 1 is an FB2N and 3 are FFNs. MBON-d1 and FAN-9 provide both ipsilateral and contralateral pathways implying these are particularly strong inputs. Input from CNs implies the existence of innate odour pathways. Input from FBNs suggest the same information used to inform memory-based action selection is also used to guide future learning. 5 of the inputs were previously published as reliable inputs to DAN-d1 (marked with *), a dopaminergic neuron that drives aversive memory likely through modulation of CS input to MBON-d1, another strong Ipsigoro input.

#### Supplementary Table 2

| No. | Abbreviated Name | Genotype | Figure | Source |
|---|---|---|---|---|
| 1 | Or42b-lexA><br>lexAop-CsChrimson | 13XLexAop2-CsChrimson-tdTomato (attP18);<br>Or42b-LexAp65 in JK22C | **3b,c** | Zlatic lab (JRC)<br>FlyStore #3025777 |
| 2 | Basin-Gal4><br>UAS-dTrpA1 | w; UAS-dTRPA1,<br>13XLexAop2-GCaMP6s 50.641 (Su(Hw)attP5);<br>72F11-GAL4 (attP2)/CyO::TM6b | **3b,c** | Zlatic lab (JRC)<br>FlyStore #3027713 |
| 4 | Goro-lexA><br>lexAop-GCaMP,<br>UAS-CsChrimson | LexAop2-Syn21-opGCaMP6s(su(Hw)attP8),<br>10XUAS-Syn21-Chrimson88-tdT-3.1(attP18);<br>69E06-LexAp65 (attp40) ; +/TM2 | **4.3a,b,<br>6.2a-d,<br>S2a** | This thesis<br>derived from stocks 16, 17 |
| 5 | empty-Split-Gal4 | w[1118]; P{p65.AD.Uw}attp40;<br>P{GAL4.DBD.Uw}attp2 | **4.3c,d,<br>5.2c,d,<br>S1a-c** | Bloomington #79603<br>(Pfeiffer et al., 2010) |
| 6 | UAS-dTrpA1 | w-; UAS-dTRPA1 | **4.3c,d,** | Derived from<br>Bloomington #26263<br>(Paul Garrity) |
| 7 | MD-IV-lexA | w[1118]; ppk-1kb-hs43-LexA-GAD (attP40);<br>MKRS/ TM6B | **5.2a,b** | Zlatic lab (JRC)<br>FlyStore #1145492<br>(Vogelstein et al., 2014) |

| | | | | |
|---|---|---|---|---|
| 8 | Ipsigoro-Gal4><br>UAS-GCaMP,<br>lexAop-CsChrimson | 13XLexAop-CSChrimson-tdTomato in attP18,<br>20XUAS-IVS-jGCaMP8s}su(Hw)attP8;<br>SS03731; +/TM6B | **5.2a,b,<br>6.1b,c,<br>S2b,c** | This thesis<br>derived from stocks 3,18 |
| 9 | MD-IV-lexA><br>lexAop-dTrpA1,<br>UAS-Shibire$^{ts1}$ | ppk-LexA (attp40);<br>pJFRC100-20XUAS-TTS-Shibire-ts1-p10 (attP2),<br>pJFRC26-13XLexAop2-IVS-dTrpA1-WPRE<br>(VK00005) | **5.2c,d** | Zlatic lab (JRC)<br>FlyStore #1145586 |
| 10 | MBON-d1-lexA | VT007911-LexA (attP40) | **6.1b,c** | Bloomington #604542<br>(Meissner et al., 2024; Tirian & Dickson, 2017) |
| 11 | MBON-d1-Gal4 | w[1118]; P{R11E07-p65.AD}attp40;<br>P{R52H01-GAL4.DBD}attp2 (SS01705) | **6.2a,b** | Bloomington #604148<br>(Eschbach et al., 2021) |
| 12 | MBON-k1-Gal4 | w[1118];<br>P{VT033301-p65.AD}attp40;<br>P{R27G01-GAL4.DBD}attp2 (SS01962) | **6.2c,d** | Bloomington #603462<br>(Saumweber et al., 2018) |
| 13 | UAS-CsChrimson | 20XUAS-CsChrimson-mVenus (attP18) | **S1a** | Bloomington #55134<br>(Klapoetke et al., 2014) |
| 14 | Basin-Gal4><br>UAS-CsChrimson | pUAS-CsChrimson-mVenus (attP18);;<br>72F11-Gal4 (attp2) | **S1b,c** | Zlatic lab (JRC)<br>FlyStore #3020667 |
| 15 | w$^{1118}$ | w[1118] | **S2a-c** | Janelia Research Campus<br>FlyStore #1500005<br>(Ryder et al., 2004) |

| | | | | |
|---|---|---|---|---|
| 16 | Goro-lexA | 69E06-LexAp65 (attp40) | | Bloomington #54925<br>(Ohyama et al., 2015; Pfeiffer et al., 2010) |
| 17 | lexAop-GCaMP,<br>UAS-CsChrimson | LexAop2-Syn21-opGCaMP6s(su(Hw)attP8),<br>10XUAS-Syn21-Chrimson88-tdT-3.1(attP18);<br>CyO/Sp;MKRS/TM2 | | Zlatic Lab (Cambridge) |
| 18 | UAS-GCaMP,<br>lexAop-CsChrimson | 13XLexAop-CSChrimson-tdTomato in attP18,<br>20XUAS-IVS-jGCaMP8s}su(Hw)attP8;<br>Sp/CyO;MKRS/TM6B | | Zlatic Lab (JRC)<br>FlyStore #3019092 |

**Supplementary Table 2. Fly Stocks used in this thesis.**
